# Xrn1 is a deNADding Enzyme Modulating Mitochondrial NAD Levels

**DOI:** 10.1101/2021.06.25.449970

**Authors:** Sunny Sharma, Jun Yang, Ewa Grudzien-Nogalska, Megerditch Kiledjian

**Affiliations:** Department of Cell Biology and Neuroscience, Rutgers University, Piscataway, NJ 08854, USA

**Keywords:** NAD cap, Xrn1, NAD cap decapping, deNADding, mitochondria

## Abstract

The existence of non-canonical nicotinamide adenine diphosphate (NAD) 5’-end capped RNAs is now well established. Nevertheless, the biological function of this nucleotide metabolite cap remains elusive. Here, we show that the yeast *Saccharomyces cerevisiae* cytoplasmic 5’-end exoribonuclease Xrn1 is also a NAD cap decapping (deNADding) enzyme that releases intact NAD and subsequently degrades the RNA. The significance of Xrn1 deNADding is evident in a deNADding deficient Xrn1 mutant that still retains its 5’-monophosphate exonuclease activity. This mutant reveals Xrn1 deNADding is necessary for normal growth on non-fermenting sugar and is involved in modulating mitochondrial NAD-capped RNA levels and in turn intramitochondrial NAD levels. Our findings uncover a functional role for mitochondrial NAD-capped RNAs as a reservoir to maintain overall NAD homeostasis. We propose NAD-capped RNAs function as a cistern for mitochondrial NAD with Xrn1 serving as a rheostat for NAD-capped RNAs.

## INTRODUCTION

Nucleotide metabolites serve as cofactors for multiple enzymatic activities essential for cellular survival and propagation ^1^. Recent demonstrations of the redox cofactor NAD covalently linked to the 5’-end of RNAs as an RNA cap in bacteria, yeast, plants and mammals has uncovered a modulatory network where a metabolite can directly impact gene expression ^2–5^. Biochemical studies in prokaryotes have shown that bacterial polymerases can add an NAD cap in place of ATP at the 5’-end during transcription initiation ^6^. On the other hand, detection of an NAD cap on post transcriptionally processed intronic small nucleolar RNAs (snoRNAs) in mammalian cells has advocated the existence of a post transcriptional NAD capping mechanism as well ^3^. These studies have also led to the identification of a battery of NAD-cap decapping (deNADding) enzymes – NudC in bacteria ^2,7,8^ and its human homolog Nudt12 ^9,10^, Nudt16 ^10^ and the DXO/Rai1 family of non-canonical decapping enzymes ^3,11^. DXO/Rai1 are class 1 deNADding enzymes that target the phosphodiester linkage and releases an intact NAD while NudC, Nudt12 and Nud16 are class 2 deNADding enzymes that hydrolyze the pyrophosphate bond within the NAD of the NAD cap and release nicotinamide mononucleotide (NMN) ^9,12^.

Functional analyses of the NAD-capped RNAs have revealed disparate roles in bacterial and mammalian cells. In bacteria, the NAD cap is considered to protect the RNA from 5’-end degradation ^2^, whereas in mammals ^3^ and plants ^13^ NAD caps have been shown to promote RNA decay. In bacteria, NAD caps are found on certain regulatory RNAs ^2^, while in budding yeast *Saccharomyces cerevisiae*, they are on a subset of mitochondrial genes and genes encoding the translational machinery ^4^. NAD-capped RNAs are prevalent in the transcriptome of plants ^5^ and human cells ^3^, and are mainly comprised of mRNAs and small nucleolar RNAs (snoRNAs). The NAD cap does not appear to support translation in mammalian cells ^3^, while their presence on plant polysomes indicates a translational capacity ^5^. Despite the occurrence of NAD caps in diverse organisms, the biological role of this non-canonical metabolite cap in cellular physiology has remained elusive.

## RESULTS

### Identification of Xrn1 and Rat1 as NAD cap associated proteins

To begin unravelling the functional role of NAD caps, we set out to identify proteins that specifically bind to the NAD cap. Protein purification was carried out by 5’**N**AD **c**ap **R**NA **A**ffinity **P**urification (NcRAP) with the RNA containing a 3’-biotin to immobilize the affinity matrix and identify proteins associated with the NAD cap from *S. cerevisiae* lysate as illustrated in Figure 1a. Canonical m^7^G-capped RNA was used as a control for the cap affinity purification and associated proteins were identified by mass spectrometry. As expected, the nuclear (Sto1 and CBC2) and cytoplasmic (eIF4E) canonical cap binding proteins were prominent bands selectively detected on the m^7^G cap RNA affinity column as well as the associated eIF4F adapter complex proteins eIF4G1 and eIF4G2 ^14^ (Figure. 1b), demonstrating the feasibility of the NcRAP approach.

**Figure 1.**
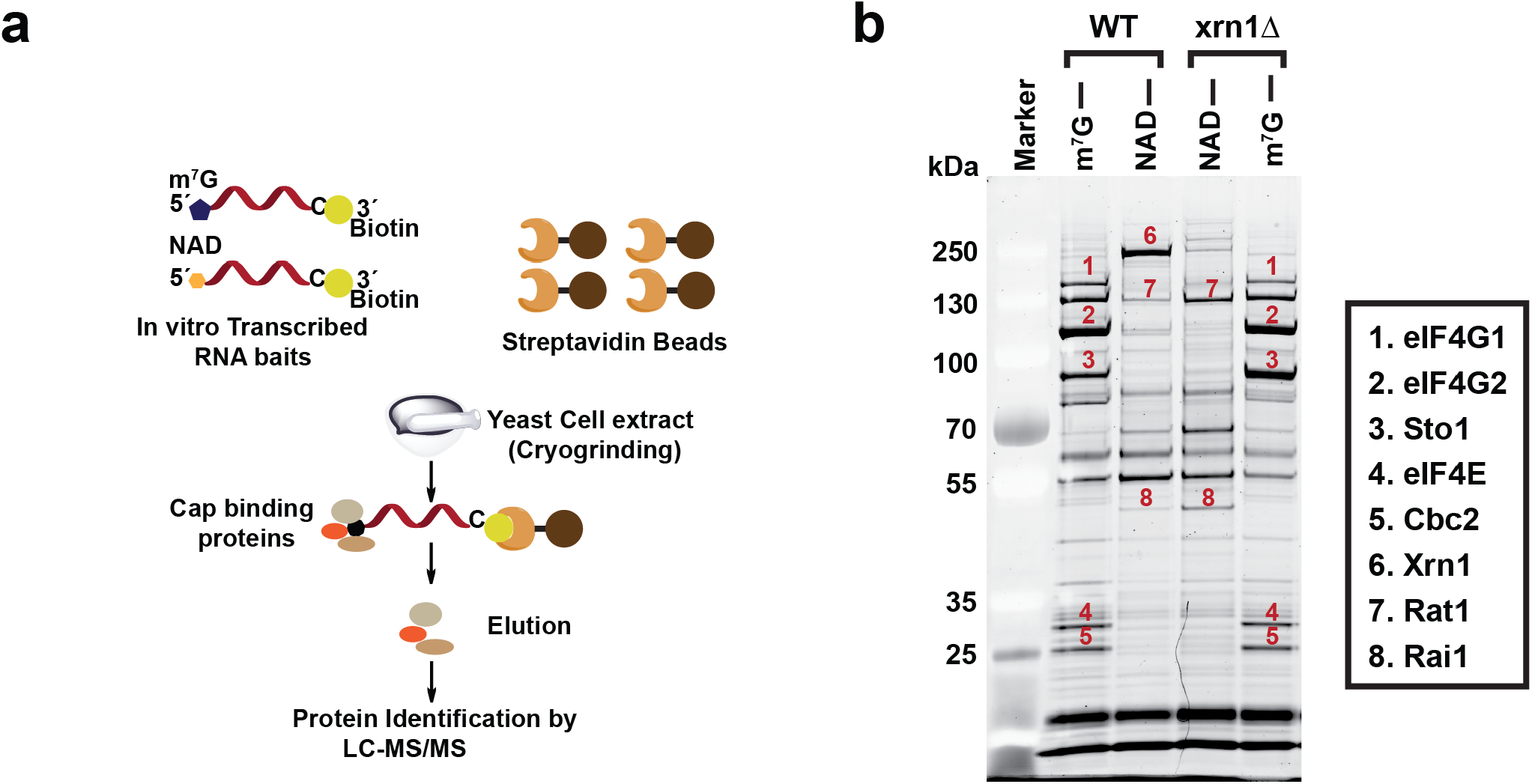
Identification of NAD cap binding proteins in budding yeast. **(a)** Schematic illustration of the NAD cap -RNA Affinity Purification (NcRAP). **(b)** Eluates from NcRAP were loaded on to a 4%-12% Bis-Tris gel and stained with SYPRO Ruby. The m^7^G cap affinity purification was used as a control. All protein bands labeled on the gel were excised from the gel and identified using mass spectrometry.

Mass spectrometry identification of proteins selectively bound to the NAD-capped RNA (Supplementary Table S1) revealed surprising candidates. The most prominent protein interacting with the NAD cap was the cytoplasmic 5’-3’ exoribonucleases Xrn1 ^15^, followed by the nuclear 5’-3’ exoribonucleases Rat1 (Xrn2 in mammals) ^16,17^ and the Rat1 interacting partner, Rai1 ^18^ (Figure. 1b). These interactions were further validated by performing NcRAP using protein lysates derived from an Xrn1 deletion (*xrn1Δ*) strain. In the absence of Xrn1, binding of both the Rat1 and Rai1 proteins to the NAD cap was accentuated. Moreover, since Rat1 and Rai1 exist as heterodimers ^18^ and Rai1 is a known deNADding protein ^3^, extract from the *rai1Δ* strain was used in NcRAP. As shown in Supplementary Figure S1, the association of Rat1 with the NAD cap was independent of its interaction with the Rai1 based on its retention onto the NAD-capped RNA column. These findings reveal Xrn1, Rat1 and Rai1 are capable of selectively binding to the NAD cap under the assay conditions employed.

### Xrn1 and Rat1 hydrolyze NAD capped RNAs *in vitro*

Identification of Xrn1 and Rat1 by NcRAP in addition to Rai1, a potent deNADding enzyme^3^, raised the intriguing possibility that analogous to Rai1, Xrn1 and Rat1 may also possess deNADding activity. Since both Xrn1 and Rat1 are highly processive 5’-monophosphate (5’P)-specific 5’-3’ exoribonucleases ^15,17^, we assessed the potential of these enzymes to hydrolyze 5’NAD-capped RNA (Figure 2a) *in vitro*. As expected, Xrn1 efficiently degraded 5’P RNA (Figure 2b) but not a 5’-triphosphate- or an m^7^G-capped RNA (Supplementary Figure S2a). Importantly, uniformly ^32^P-labeled NAD-capped RNA was degraded by wild type *Kluyveromyces lactis* (Figure 2b) and *S.cerevisiae* (Figure S2b) Xrn1, but not the catalytically inactive *xrn1-E178Q* ^19^ mutant protein. Interestingly, the decay activity of Xrn1 on NAD-capped RNA was processive without detectable intermediates in contrast to *Schizosaccharomyces pombe* Rai1, which removed the NAD cap without subsequent degradation of the deNADded RNA (Figure 2b), consistent with our previous reports ^3^. Similarly, *S.pombe* Rat1 also hydrolyzed the NAD-capped RNA with processive exonuclease activity, whereas the catalytically dead *rat1-E207Q* ^20^ was inactive (Supplementary Figure S2c). Collectively, these data demonstrate that the observed degradation of the NAD-capped RNA is a function of Xrn1 and Rat1, and requires the same catalytic active site utilized for the hydrolysis of a 5’P RNA.

**Figure 2.**
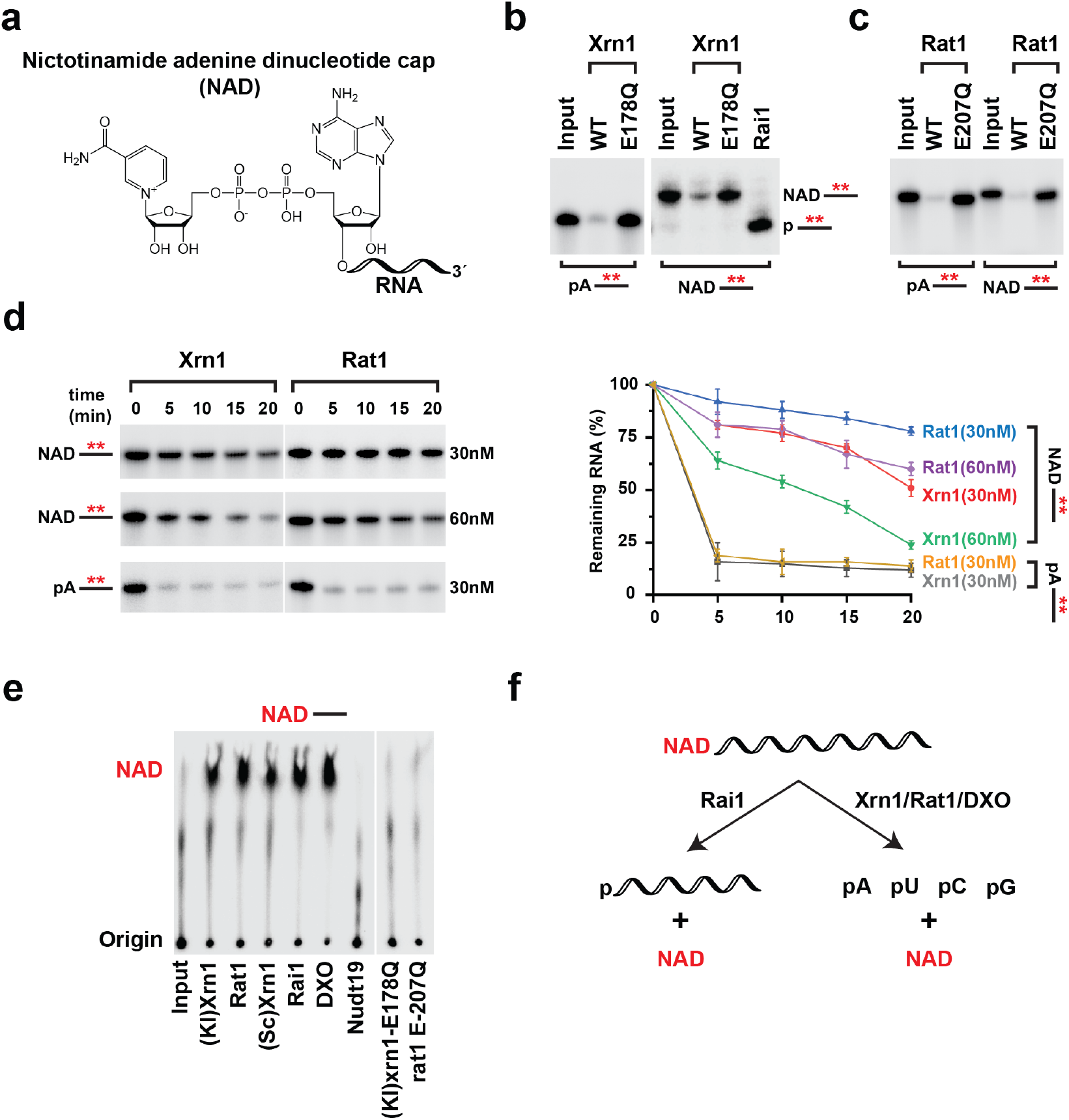
Xrn1 and Rat1 are deNADding enzymes. **(a)** Structure of NAD-capped RNA. **(b)** Reaction products of *in vitro* deNADding assays with 30 nM recombinant Xrn1, WT, and catalytically inactive (*E178Q*) from *K. lactis or* 25 nM Rai1 (*S. pombe*), **(c)** and 50 nM recombinant Rat1, WT or catalytically inactive (*E207Q*) from *S. pombe*. Uniformly ^32^P-labeled monophosphate (pA----) or NAD-capped RNA (NAD---) where the line denotes the RNA were used as indicated. The red asterisk represents the ^32^P-labeling within the body of the RNA. Reaction products were resolved on 15% 7M urea PAGE gels with **(b)** and without **(v)** 0.3% 3-acrylamidophenylboronic acid. **(d)** Time-course decay analysis of uniformly ^32^P-labeled monophosphate or NAD-capped RNA with the indicated amount of Xrn1 or Rat1 protein are shown. Quantitation of RNA remaining is plotted from three independent experiments with ± SEM denoted by error bars. **(e)** NAD-capped RNA containing the ^32^P-labelling within the a phosphate of the NAD (labeled in red) was subjected to the indicated proteins Decay products were resolved by polyethyleneimine (PEI)-cellulose thin layer chromatography (TLC) developed in 0.45M (NH4)2SO4. Rai1 and DXO served as positive controls, whereas Nudt19 was a negative control. **(f)** Schematic of NAD-capped RNA hydrolysis by Rai1, Xrn1, Rat1 and DXO.

To further understand the kinetics of deNADding, a time-course decay assay was carried out with 5’P- or NAD-capped RNA using two different concentrations of Xrn1 and Rat1. Consistent with the well characterized exoribonuclease activities of Xrn1 and Rat1, 5’P RNA was degraded more efficiently than NAD-capped RNA at comparable enzyme concentrations (Figure 2d). Furthermore, Xrn1 degraded the NAD-capped RNA twice as efficiently as Rat1. To assess whether Xrn1 hydrolyzed the NAD cap within the diphosphate moiety analogous to the class 2 Nudt12 protein, or removed the intact NAD, similar to the class 1 DXO/Rai1 family of proteins ^12^, thin layer chromatography (TLC) was used. Interestingly, ^32^P-labeled NAD-capped RNA subjected to Xrn1 and Rat1 hydrolysis released intact NAD (Figure 2e). In addition, unlike Nudt12 both Xrn1 and Rat1 do not hydrolyze the diphosphate bond (Supplementary Figure S2d). We conclude that Xrn1 and Rat1 are class 1 deNADding enzymes that have the capacity to degrade NAD-capped RNA analogous to the DXO/Rai1 family of proteins by removing the intact NAD (Figure 2f). These findings increase the repertoire of substrate RNAs regulated by Xrn1 and Rat1 beyond their well characterized 5’-monophosphate directed exoribonuclease activity.

### Loss of Xrn1 stabilizes NAD-capped RNAs *in vitro* and *in vivo*

Having demonstrated Xrn1 and Rat1 function as deNADding enzymes *in vitro*, we assessed the deNADding activity of Xrn1 in cells. We focused on Xrn1 since the *xrn1Δ* strain is viable and amendable to genetic modifications while the *rat1Δ* stain is not ^16^. 5’-end RNA decay assays were carried out with extract derived from either wild-type (WT) or *xrn1Δ* cells with NAD-capped or m^7^G-capped RNAs immobilized and protected at the 3’-end with biotin-streptavidin (Figure 3a). While no significant differences were observed with the control m^7^G-capped RNA, the NAD-capped RNA was more stable in the *xrn1Δ* extract, suggesting endogenous Xrn1 possesses deNADding activity (Figure 3b). To assess the contribution of Xrn1 deNADding on endogenous NAD-capped RNA, we used the NAD cap detection and quantitation (NAD-capQ) approach that detects NAD caps released from the 5’-end of RNAs *en masse* (Supplementary Figure S3a) ^21^. Consistent with a deNADding function for Xrn1 in cells, a statistically significant 1.5-fold higher level of total NAD cap was detected in the *xrn1Δ* strain relative to the WT strain (Figure 3c).

**Figure 3.**
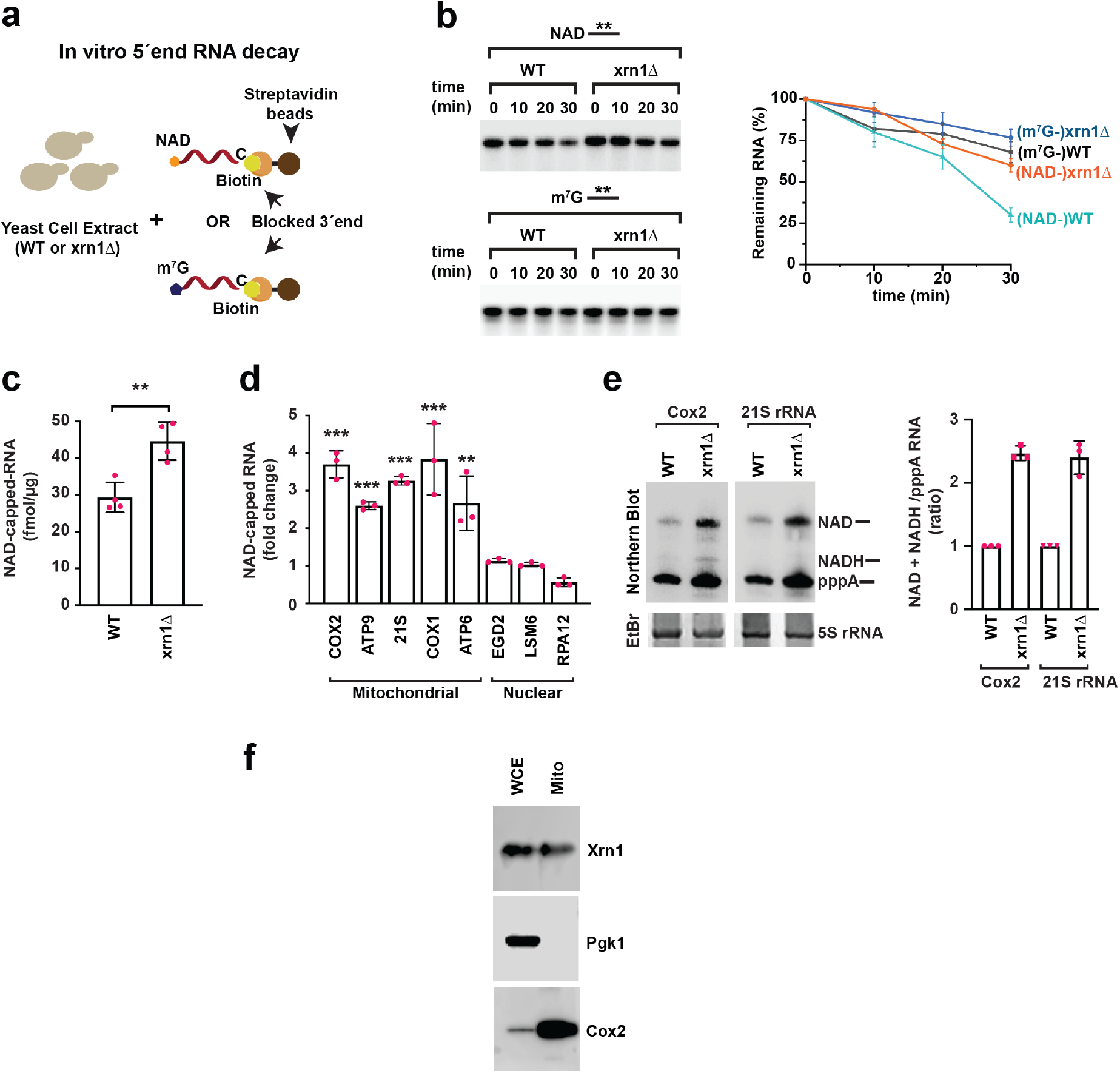
Xrn1 deNADs NAD-capped RNAs in cells. **(a)** Schematic illustration of in vitro 5’-end RNA decay. **(b)** Time-course decay analysis of uniformly ^32^P-labeled m^7^G- or NAD-capped RNA in the presence of cell extracts prepared from WT or xrn1Δ strains. The remaining RNA was quantified and plotted from three independent experiments with ± SEM denoted by error bars. Labeling as in the legend to Figure. 2. **(c)** Total RNAs from WT and xrn1Δ were subjected to the NAD-capQ assay to detect total level of cellular NAD-capped RNAs. Data represents average from three independent experiments. Error bars represent ± SEM. (Average of three independent experiments with error bars represent ± SEMmean ± SEM; unpaired t-test; * p < 0.05, ** p < 0.001, *** p < 0.0001). **(d)** qRT–PCR quantification of NAD-capped mRNAs in WT and xrn1Δ strains. NAD-capped RNAs were enriched by NAD-capture, eluted from the beads, reverse transcribed and detected with gene-specific primers. Data are presented relative to the WT cells set to 1 and represent three 3 independent NAD-capture experiments; mean ± SEM; unpaired t-test; * p < 0.05, ** p < 0.001, *** p < 0.0001. **(e)** Northern Blot analysis of DNAzyme-generated 5’-end-containing subfragments of Cox2 and 21S rRNA with the ^32^P labeled transcript specific probes are shown with the corresponding quantitation plotted on the right. Data are derived from three independent experiments with error bars represent ± SEM. Schematics of the NAD-capQ and DNAzyme approaches are shown in Supplementary Figure S3. **(f)** Density gradient centrifugation purified mitochondria derived from a strain harboring Xrn1 with a C-terminus Strep-tag II was analyzed by Western blot analysis with an anti-Strep-tag antibody. Western blotting analysis of whole cell extract (WCE) and extract prepared from purified mitochondria (mito) using Pgk1 (cytoplasmic protein) and Cox2 (mitochondrial protein) were used to determine the purity of mitochondria.

### Xrn1 deNADding activity influences NAD-capped mitochondrial transcripts

Previous transcriptome-wide analyses of NAD-capped RNA in budding yeast identified distinct NAD-capped nuclear and mitochondrial transcripts ^4,22^ with levels as high as 50% of select mitochondrial RNAs containing an NAD cap ^22^. To determine whether accumulation of endogenous NAD-capped RNAs were altered in the absence of Xrn1, NAD-capped RNAs were isolated by NAD capture ^23^ and subjected to RT-qPCR analyses. We focused on a subset of previously identified NAD-capped nuclear- and mitochondrial encoded mRNAs ^4^. Importantly, although loss of *xrn1* did not change the steady state accumulation of the nuclear encoded NAD-capped RNAs tested, all the mitochondrially encoded NAD-capped transcripts tested were significantly elevated (Figure 3d). Furthermore, we validated the qPCR results by an independent approach using boronate affinity electrophoresis for two transcripts, Cox2 and 21S rRNA ^22^. This method allows visualization of distinct capped and uncapped RNA populations by specifically retarding the mobility of NAD-capped RNAs in the gel due to transient formation of diesters between immobilized boronic acid and the nicotinamide moiety ^22,24^. DNAzyme-mediated RNA cleavage was used to generate 5’-end containing sub fragments of defined length (Supplementary Figure S3b). Northern blot analysis with the ^32^P labeled transcript-specific probes corroborated the NAD capture analyses for Cox2 and 21S rRNA with elevated NAD-capped RNA in the *xrn1Δ* strain (Figure 3e). This observation prompted us to test whether Xrn1 is associated with mitochondria. In addition to its well characterized cytoplasmic localization ^25^, Xrn1 has also been reported on cell membrane compartments called eisosomes that mark the sites of endocytosis during post-diauxic shift ^26^. Gradient centrifugation purified mitochondria derived from a strain harboring Xrn1 with a C-terminus Strep-tag II was analyzed by Western blot analysis with an anti-Strep-tag antibody. As shown in figure 3f, Xrn1 was detected with mitochondrial preparations suggesting Xrn1 can be mitochondrially associated. Together, both the *in vitro* reconstitution and the *in vivo* analysis demonstrated a functional role for Xrn1 in regulating the fate of NAD-capped RNA in yeast mitochondria.

### A conserved histidine residue in the catalytic pocket of Xrn1 is indispensable for NAD-capped RNA hydrolysis

Structural and functional analyses of Xrn1 from *K. lactis* and *Drosophila melanogaster* have elucidated the molecular basis of the 5’P-specificity and the mechanism of processive RNA degradation ^19,27^. The 5’P is embedded in a highly basic pocket lined with the side chains of the conserved amino acids K93, E97, R100, and R101 with His41 providing directionality to the decay process ^27^ (Supplementary Figure S4a and S4b). To determine whether the 5’P hydrolysis activity of Xrn1 can be uncoupled from the deNADding activity, a mutational screen was initiated. Structure guided point mutations were generated in the 5’P binding pocket and the catalytic residues and assessed for their deNADding activity. Importantly, one residue, His41, was indispensable for deNADding while it still supported 5’P RNA decay activity (Figure 4a, Supplementary Figure S4c). To gauge the impact of H41A on deNADding in cells, a *xrn1-H41A* knock-in mutant strain was generated. Consistent with the *in vitro* analysis, the H41A mutant retained 5’P mediated exonuclease activity in cells as shown by its capacity to process its natural nuclear encoded substrates, Rpl22 precursor RNA ^28^ (Figure 4b) and Rpl28 precursor RNA ^29,30^ (Supplementary Figure S5). We next tested the consequence of a deNADding deficient Xrn1 on NAD-capped RNA levels in cells and detected an elevation of total NAD-capped RNA in the *xrn1-H41A* knock-in strain (Figure 4c). Moreover, direct detection of the Cox2 and 21S rRNA mitochondrial transcripts revealed a four- and three-fold elevation of NAD-capped RNA in the *xrn1-H41A* strain respectively (Figure 4d). We conclude that H41 is a critical residue for the Xrn1-directed deNADding activity in cells and provides an avenue to directly determine the functional significance of Xrn1 deNADding in cells.

**Figure 4.**
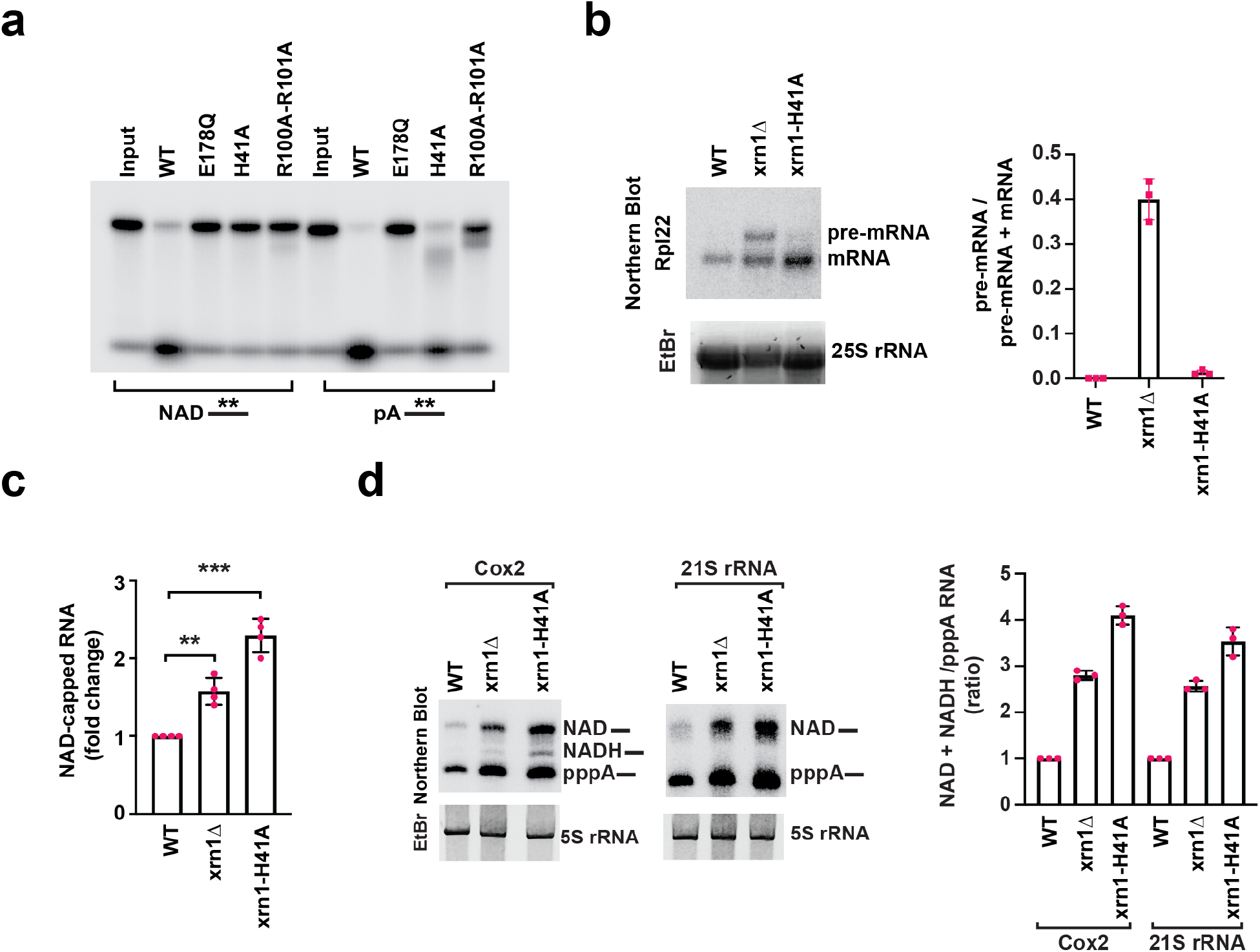
A conserved histidine residue in the catalytic pocket of Xrn1 is indispensable for NAD capped RNA hydrolysis. Xrn1 mutational screen revealed that H41 is indispensable for NAD-capped RNA hydrolysis. **(a)** 30 nM recombinant WT or mutated Xrn1 from *K. lactis* was incubated with uniformly ^32^P-labeled 5’-monophosphate or NAD-capped RNA. The products were resolved on 15% 7M urea PAGE gels. **(b)** Northern Blot for Rpl22 mRNA in WT, *xrn1Δ* and *xrn1*-H41A knock-in mutant. Quantitation from three independent experiments are presented on the right with the mean ratio of Rpl22 pre-mRNA over total (pre-mRNA + mRNA). **(c)** Total RNAs from WT, *xrn1Δ, xrn1*-H41A strains were subjected to the NAD-capQ assay to detect total level of cellular NAD-capped RNAs. Average from three independent experiments are shown with error bars denoting ± SEM. Statistical significance level was calculated by one-way ANOVA (F = 18.30, df = 2.209, p < 0.0001) with a Dunnett’s multiple comparison post hoc test, * p < 0.05, ** p < 0.001, *** p < 0.0001. **(d)** Northern Blot analysis of DNAzyme-generated 5’-end-containing subfragments of Cox2 and 21S rRNA from WT, *xrn1Δ* and *xrn1*-H41A grown in YPD (fermentation) media. Quantitation from three independent experiments with error bars representing ± SEM are shown on the right.

### Xrn1 maintains NAD-capped RNA and mitochondrial NAD level homeostasis

The 5’-3’ exonuclease activity of Xrn1 is known to be necessary for growth on nonfermentable carbon sources in yeast resulting in a slow growth phenotype in its absence ^31^. We next determined whether Xrn1 deNADding activity contributed to the inhibitory growth on non-fermenting sugar utilizing the deNADding-deficient *xrn1-H41A* strain. As expected, all strains exhibited commensurate growth under fermentation growth parameters with glucose which would not require mitochondrial activity for growth (Figure 5a). Interestingly, the deNADding compromised *xrn1-H41A* strain demonstrated a slow growth phenotype comparable to the *xrn1* deletion strain on glycerol as the non-fermenting carbon source (Figure 5a) where mitochondrial activity is required for cellular energetics. Importantly, a corresponding increase in NAD capped RNA was also evident in *xrn1-H41A* mutant mitochondrial RNA (Figure 5b), indicating a correlation between NAD capped RNA and cellular growth when mitochondrial function was essential.

**Figure 5.**
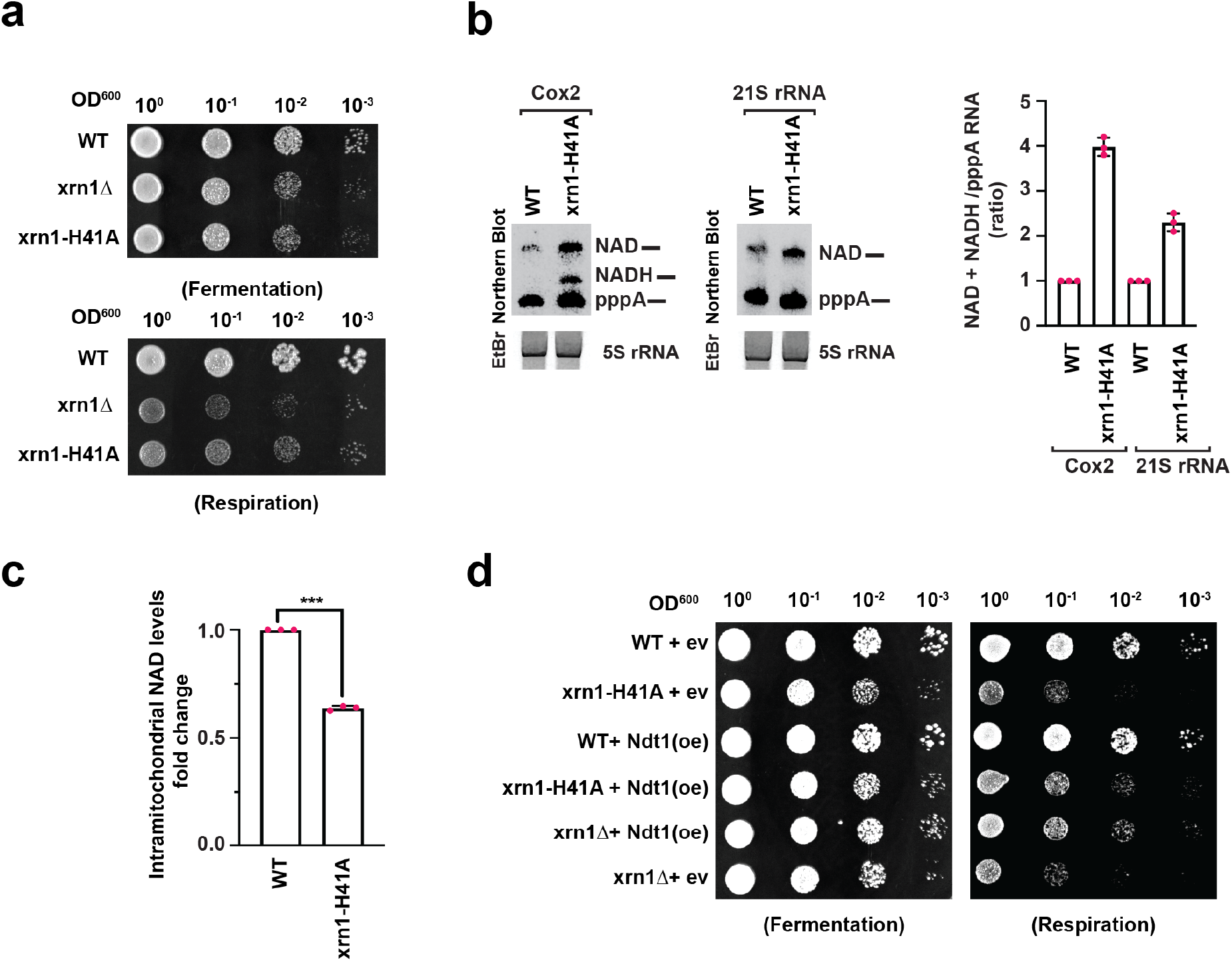
Xrn1 deNADding activity is critical in maintaining intramitochondrial NAD steady state levels. **(a)** Tenfold serial dilution of WT, *xrn1Δ* and *xrn1*-H41A were spotted onto solid YPD (Fermentation) and YPG (Respiration) plates and incubated at 30°C. **(b)** Northern Blot analysis of DNAzyme-generated 5’-end-containing subfragments of Cox2 and 21S rRNA from WT, *xrn1Δ* and *xrn1*-H41A grown in YPG (respiration) media with the quantitation on the right (Average of three independent experiments with error bars represent ± SEM mean ± SEM). **(c)** Intramitochondrial NAD/NADH levels in WT and *xrn1*-H41A cells grown on YPG. Data are presented relative to the WT cells set to 1 and derived from three biological replicates (Average of three independent experiments with error bars represent ± SEM mean ± SEM; unpaired *t*-test; * p < 0.05, ** p < 0.001, *** p < 0.0001). **(d)** Tenfold serial dilution of WT, *xrn1Δ* and *xrn1*-H41A harboring either an empty vector (ev) – without Ndt1 or a multicopy plasmid containing Ndt1 under a strong promoter – over expressed (oe) were spotted onto solid YPD (Fermentation) and YPG (Respiration) plates and incubated at 30°C.

To determine whether levels of NAD-capped RNA influenced overall mitochondrial NAD levels, NAD/H levels were determined from WT and *xrn1-H41;A* mitochondria derived under respiring growth conditions. Our findings revealed a ~50% reduction of intramitochondrial NAD levels in the *xrn1-H41A* mutant strain (Figure 5c). With NAD capped RNAs constituting a significant fraction of mitochondrial RNA in yeast ^22^, these data imply NAD-capped RNAs can function as a cistern for mitochondrial NAD with Xrn1 serving to impact the release of the NAD from RNAs to maintain proper homeostasis. To further validate that the observed slow growth phenotype of the *xrn1-H41A* strain was a consequence of reduced mitochondrial NAD levels, we overexpressed the main mitochondrial NAD transporter protein NDT1, which leads to elevated intramitochondrial NAD ^32^. As shown in Figure 5d, overexpressing of NDT1 reversed the slow growth phenotype observed in both the *xrn1Δ* and *xrn1-H41A* strains demonstrating the slow growth phenotype is a function of insufficient free NAD in the mitochondria. We conclude Xrn1 regulates NAD homeostasis in the mitochondria through its deNADding activity and may link RNA metabolism to more general mitochondrial health and potential mitochondriopathies.

## DISCUSSION

Pyridine dinucleotides, particularly nicotinamide adenine dinucleotide (NAD) and its reduced form NADH are essential players in a variety of cellular processes, including energy metabolism, cellular signaling, and transcription^1^. Recent mass spectrometry^23,33,34^ and chemoenzymatic analysis ^23^ of cellular transcripts in organisms representing all three kingdoms of life have revealed that these dinucleotides also serve as non-canonical 5’-end RNA caps ^12^. Nevertheless, the precise role of these non-canonical caps remained unknown.

In the present study, we characterized cellular proteins from the yeast *S.cerevisiae* interacting with a 5’NAD cap using NAD cap RNA affinity purification (NcRAP) and mass spectrometry. Our data revealed that unlike the canonical m^7^G caps, the NAD cap do not interact with translational factors or proteins involved in cellular translation, suggesting that NAD-capped RNAs are unlikely to be translated in budding yeast consistent, with the lack of yeast extract supporting NAD capped RNA translation *in vitro* ^35^. Surprisingly, herein we revealed that highly conserved 5’-monophosphate (5’P)-specific 5’-3’ exoribonucleases, Xrn1 and Rat1 along with its interacting partner Rai1 (yeast homolog of human DXO) can associate with and hydrolyze 5’NAD-RNA *in vitro*. Notably, similar to the DXO/Rai1 family of proteins, both Xrn1 and Rat1 release intact NAD from the NAD-capped RNA and have the capacity to regenerate NAD.

Another unexpected outcome of our study was the influence of Xrn1 on NAD-capped mitochondrial RNAs. At present our data indicates a clear mitochondrial association of Xrn1 by its presence in highly purified mitochondrial fraction. It is possible that its deNADding activity in the proximity of the mitochondria may indicate membrane association analogous to its localization with eisosomes ^26^, and indirectly influence mitochondrial RNA deNADding. A more provocative explanation would be an intra mitochondrial localization of this well-established cytoplasmic protein to directly exert a function in the mitochondria. More detailed localization studies will be necessary to distinguish these two possibilities.

Although our initial objective was to identify NAD cap-binding proteins, the prominent proteins associated with the NAD cap were RNA nucleases and not *bona fide* NAD cap-binding proteins. This finding is consistent with our previous report that the NAD cap functions as a tag to promote rapid RNA decay ^3^. It is possible that eukaryotic cells are devoid of NAD cap-binding proteins and NAD-capped RNAs instead are recognized by RNA nucleases to rapidly clear the RNA. More refined attempts with extract derived from a stain lacking the three dominant NAD cap associated proteins, Xrn1, Rai1 and a conditional deletion of Rat1, may be required to detect additional bound proteins. In addition to Xrn1, we also demonstrated that Rat1 is capable of binding and hydrolyzing NAD-capped RNAs. Since Rat1 and Rai1 are primarily nuclear proteins ^25^ it is likely that their activity is predominantly on nuclear NAD-capped RNAs distinct from Xrn1. A more holistic and comparative analysis of the NAD-capped RNAs in the mutant strains of these three deNADding enzyme will aid in understanding their distinct or overlapping NAD-capped RNA substrates.

Mutational analysis of the catalytic center of Xrn1 led to the identification of a conserved histidine residue – His41, which is indispensable for deNADding, but does not significantly affect 5’P hydrolysis. Importantly, this residue allowed for the uncoupling of the deNADding activity of Xrn1 from the 5’P exoribonuclease activity and enabled a delineation of the physiological role of NAD capped RNA in yeast. Moreover, based on the observations of the *xrn1-H41A* knock-in mutant cells, we suggest that the previously characterized slow growth phenotype of *xrn1Δ* on non-fermenting sugar ^31^ is likely due to the loss of deNADding activity and not its monophosphate directed RNA decay.

The intracellular concentration of pyridine dinucleotides is tightly regulated in cells ^36^, therefore we hypothesized that an increase in the NAD-capped RNA or incorporation of NAD in RNA would lead to a decrease in intramitochondrial NAD levels. Consistent with this hypothesis, we observe ~50% reduction in the intramitochondrial NAD levels in *xrn1-H41A* strains grown in YPG (non-fermenting sugar). Since mitochondria are indispensable for respiration, a decrease in NAD levels in mitochondria result in a significant slow growth phenotype upon growth on YPG media. It is important to note that mitochondria of budding yeast lack enzymes necessary for NAD synthesis and primarily rely on NAD imported from the cytoplasm ^37^. In addition, the availability of free NAD dictates levels of mitochondrial NAD-capped RNA synthesis ^22^. We propose that NAD-capped RNAs serve as a reservoir for this critical nucleotide metabolite in mitochondria and Xrn1-mediated turnover of NAD-capped RNA is vital for mitochondrial NAD homeostasis and cellular growth during respiration in budding yeast. A recent report in *Arabidopsis thaliana* reinforces the equilibrium between NAD caps and free NAD where stabilization of NAD caps leads to a decrease in free NAD and compensatory increase in NAD production following stress ^13^. Whether NAD molecules stored in the 5’ end of RNA are used in adverse scenarios like nutrient starvation or quiescence in mammalian cells and the contribution of Xrn1 in NAD homeostasis will be an important area of future exploration.

## ACKNOWLEDGEMENTS

We thank Liang Tong (Columbia Univ.) for helpful discussions, and grateful to Kiledjian lab members for their input throughout the project and critical reading of the manuscript. The research was supported by NIH grant GM126488 to MK.

## AUTHOR CONTRIBUTIONS

M.K., S.S. designed the experiments. S.S, J.Y and E.G.N carried out the experiments. M.K. and S.S wrote the manuscript with input from all coauthors.

## DECLRATION OF INTERESTS

The authors declare no competing financial interests: details are available in the online version of the paper. Correspondence and requests for materials should be addressed to M.K.

## METHODS

### Yeast growth and Media

All *Saccharomyces cerevisiae* strains used in the present study were derived in the BY4741 strain background and their genotypes are listed in Supplementary Table S3. Yeast cells were grown at 30 °C either in YPD (Dextrose) or YPG (Glycerol) (1% w/v yeast extract, 2% w/v peptone, 2% w/v glucose/glycerol). For tenfold serial dilution growth assays, yeast cells were grown overnight in YPD or YPG medium and diluted first to an OD600 of 1 and then serially 1 to 10. From each diluted cultures, 5μL was spotted onto a YPD or YPG plate and incubated at 30 °C.

### Plasmids construction and PCR mediated site-directed Mutagenesis

Polymerase chain reaction (PCR) amplification was conducted from the genomic DNA of *K. lactis* using *XRN1* gene specific primers listed in the Table S2. The PCR product was gel purified and ligated into a pET-28 vector using In-Fusion^®^ (Takara) cloning.

PCR mediated site directed mutagenesis was used for introducing specific point mutations in the Xrn1 protein. PCR with *Single-Primer Reactions IN Parallel* (SPRINP) ^38^ and high-fidelity-DNA polymerase was used. All the primers used for generating point mutations are listed in the Supplementary Table S2.

### *In vivo* site directed mutagenesis using *Delitto Perfertto*

Mutation substitution in the *XRN1* gene was generated using *Delitto Perfetto* ^39^. Briefly, a CORE (**CO**unterselectable marker and **Re**porter gene) – *URA3-kanMx* was introduced into the Xrn1 locus proximal to His40 using 40-nts flanking sequences for homologous recombination. In the second step, a PCR-amplified fragment containing the mutation of interest was transformed into the strain to enable integration and excise CORE cassette in the process. All point mutations were validated by DNA sequencing.

### Protein Expression and Purification

Recombinant proteins were expressed and purified using Nickel-NTA affinity purification. In brief, the ligated plasmid was transformed into chemical competent *E. coli* BL21 (DE3) cells by heat shock. 5 mL LB cultures containing 50 μg/mL kanamycin were inoculated with single colonies and grown overnight. These cultures were used to inoculate 1 L of the same medium and allowed to grow at 37 °C until an OD600 of 0.5. Protein expression was induced with 0.25 mM isopropyl D-thiogalactopyranoside (IPTG) and cells grown at 18 °C on a shaker for ~18 h. Cells were collected by centrifugation, 5000 g for 15 minutes.

For protein purification, cell pellets were thawed and resuspended in ~60 mL of buffer [25 mM HEPES (pH 7.5), 250 mM KCl, 10% glycerol, and 100 μM MnCl2] containing 0.4 mg/mL protease inhibitor phenylmethanesulfonylfluoride fluoride (PMSF). Cells were lysed by sonication, and the insoluble cell debris was separated by centrifugation. The Nickel-NTA affinity purification was performed as described previously ^40^.

### *In vitro* transcription of NAD-capped RNAs

RNAs containing NAD and m^7^G cap structures were synthesized by *in vitro* transcription from a synthetic double stranded DNA template ɸ2.5-NAD-40 containing the T7 ɸ2.5 promoter and a single adenosine within the transcript contained at the transcription start site (Supplementary Table S2) ^10^. For m^7^G-capped RNA, m^7^G(5’)ppp(5’) A RNA Cap Structure Analog (New England Biolabs) was included in the transcription reaction. *In vitro* transcription was carried out at 37 °C overnight, using HiScribe™ T7 High yield RNA Synthesis kit (New England Biolabs).

To generate ^32^P labelled NAD-capped RNA, transcription was carried out in the presence of ^32^P-NAD (PerkinElmer) instead of ATP using ɸ2.5-AG-30 (Supplementary Table S2) ^10^. To generate ^32^P labeled-RNA, the transcription reactions were performed in the presence of [α-^32^P] GTP.

### NAD cap RNA Affinity Purification (NcRAP)

400 pmol of both NAD and m^7^G capped RNAs were immobilized onto Dynabeads™ MyOne™ Streptavidin T1 (Invitrogen™) following manufacturer’s instructions. For NcRAP, yeast cells were grown in YPD at 30 °C. 2 L of exponentially grown yeast cultures were harvested and lysed using mortar and pestle in liquid nitrogen. The cell powder obtained after grinding was resuspended in lysis buffer (25 mM Tris-HCl pH 7.5, 150 mM CH3CO2K, 2.5 mM Mg(C2H3O2)2, 50 μM EDTA, 50 μM EGTA, 2.5% glycerol and 0.25% IPEGAL along with complete EDTA-free Protease Inhibitor Cocktail (Roche) and incubated at 4 °C for 30 minutes. Samples were subsequently centrifuged for 10 minutes at 5,000 g to remove the cell debris. The clarified protein lysate was then transferred to a fresh 15 mL Falcon tube and the protein concertation was determined using a Bradford assay. 5 mg of protein lysate was added to the preequilibrated (in 1X lysis buffer) NAD- or m^7^G-capped RNA coupled Dynabeads in 2mL microcentrifuge tubes. These tubes were next incubated at 4 °C on an end to end rocker for 90 minutes. The beads were washed for at least 5 times with 1X lysis buffer, and the bound proteins were eluted with 2X Laemmli buffer (Bio-Rad Laboratories). The samples were incubated at 85 °C for 5 minutes and were next run on Novex™ WedgeWell™ 4 to 20 %, Tris-Glycine, Protein Gel (Thermo Fischer Scientific). The protein gel was stained with SYPRO Ruby (Bio-Rad Laboratories) as per manufacturer’s instructions and specific bands were sliced and sent for mass spectrometry based identification. In parallel we also analyzed the entire eluate using mass spectrometry. Complete list of identified proteins are provided in the Supplementary Table S1. Mass spectrometry was carried out at Biological Mass Spectrometry Facility of Robert Wood Johnson Medical School and Rutgers, The State University of New Jersey.

### RNA *in vitro* deNADding assay

The ^32^P-NAD-cap labeled and [α-^32^P]GTP uniformly labelled NAD-capped RNAs were incubated with the indicated amounts of recombinant protein in NEB buffer 3 (100 mM NaCl, 50 mM Tris-HCl, 10 mM MgCl2, pH 7.9). Reactions were stopped by heating at 65 °C for 5 minutes. Decapping products were resolved by polyethyleneimine (PEI)-cellulose thin-layer chromatography (TLC) (Sigma-Aldrich) and developed in 0.45 M (NH4)2SO4.

For *in vitro* 5’-end RNA decay, cell extract corresponding to 2 μg of cellular protein from WT or *xrn1Δ* strains were incubated with [α-^32^P]GTP uniformly labelled NAD and m^7^G capped RNAs immobilized onto Dynabeads™ MyOne™ Streptavidin T1 (Invitrogen™) in NEB buffer 3 (100 mM NaCl, 50 mM Tris-HCl, 10 mM MgCl2, pH 7.9).

### NAD-cap detection and quantitation (NAD-capQ)

Yeast strains were grown in YPD media at 30 °C according to standard protocols. All strains were harvested for the experiments at exponential phase (OD600 ~1) and total RNA was isolated using the acidic hot phenol method ^41^. To remove residual free NAD, purified RNAs were dissolved in 10 mM Tris-HCl (pH 7.5) containing 2 M urea. Samples were incubated 2 min at 65 °C and immediately precipitated with isopropanol in the presence of 2 M ammonium acetate. NAD-capQ was carried out as previously described ^21^. Briefly, 50 μg of total RNA was digested with 2 U of Nuclease P1 (Sigma-Aldrich) in 20 μL of 10 mM Tris (pH 7.0), 20 μM ZnCl2 at 37 °C for 1h to release 5’-end NAD. The control samples lacking Nuclease P1 were prepared by incubating 50 μg of RNA treated with the same reaction condition. The NADH standard curve was generated for each experiment in the same buffer condition as above for assays containing nuclease P1. Following digestion with nuclease P1, 30 μL of NAD/NADH Extraction Buffer (NAD/NADH Quantitation Kit, Sigma-Aldrich) was added to each sample. In the second step, 50 μL samples were used in a colorimetric assay according to the manufacturer’s protocol (NAD/NADH Quantitation Kit, Sigma-Aldrich) as described ^21^.

### Isolation of NAD-capped RNAs by NAD capture and RNA quantitation

Total RNA from yeast strains grown in YPD media was isolated with the acidic hot phenol method ^41^ and treated with DNase (Promega) according to the manufacturers’ protocols. NAD-capped RNAs were isolated using the NAD-RNA capture protocol ^23^ with minor modifications. NAD-capture was carried out with 50 μg total RNA treated with 10 μL 4-pentyn-1-ol (Sigma-Aldrich) and 3 U Adenosine diphosphate-ribosylcyclase (ADPRC) in 100 μL reaction containing 50 mM HEPES, 5 mM MgCl2 (pH 7) and 40 U RNasin^®^ Ribonuclease Inhibitor (Promega) at 37° C 60 min. The reaction was stopped with phenol/chloroform extraction and RNAs were precipitated with ethanol in the presence of 2 M ammonium acetate. RNAs with a 5’-end NAD were biotinylated by treatment with a copper-catalyzed azide-alkyne cycloaddition (CuAAC) reaction by incubating with 250 μM biotin-PEG3-azide, freshly mixed 1 mM CuSO4, 0.5 mM THPTA, 2 mM Sodium Ascorbate in 100 μL reaction with 50 mM HEPES, 5 mM MgCl2 (pH 7) and 40 U RNasin^®^ Ribonuclease Inhibitor (Promega) at 30° C for 30 min. RNA was precipitated with ethanol in the presence of 2 mM EDTA and 2 M ammonium acetate. Biotinylated NAD capped RNAs were captured by binding to 20 μL streptavidin magnet beads (Nvigen) at room temperature with gentle rocking for 1 hour in 100 μL binding buffer (1 M NaCl, 10 mM HEPES (pH 7.5) and 5 mM EDTA). The beads were washed three times with buffer containing 8 mM Urea, 50 mM Tris-HCl (pH 7.4), 0.1% IPEGAL). To elute biotinylated NAD-capped RNAs, beads were resuspended in 100 μL of above buffer, heated to 95 °C for 5 min and RNA precipitated with ethanol in the presence of glycogen and 2 M ammonium acetate. Captured NAD-RNAs were dissolved in 20 μL H2O and reverse transcribed with M-MLV reverse transcriptase, random hexamers, and oligo (dT) (Promega) according to the manufacturer’s instructions. qRT-PCR was performed with the primers listed in Supplementary Table S2. qRT-PCR was carried out on QuantStudio 3 Real-Time PCR System (Thermo Fisher Scientific) with iTaq SYBR Green Supermix (Bio-Rad Laboratories).

### Boronate affinity electrophoresis and Northern Blotting analysis of *in vivo* NAD capped transcripts after DNAzyme mediated RNA cleavage

For analysis of NAD and NADH-capping during fermentation, yeast strains were grown in YPD and for respiration in YPG media at 30 °C. RNA was isolated from cells at the exponential phase (OD600 ~1) with the acidic hot phenol method ^41^ and treated with DNase (Promega). 50 μg of total cellular RNA was incubated with 1 μM of the corresponding DNAzyme ((Supplementary Table S2) in 50 μL reaction containing 10 mM Tris pH = 8.0, 50 mM NaCl, 2 mM DTT and 10 mM MgCl2. Samples were heated at 85 °C for 2 min, cool to 37 °C. MgCl2 was added to final concentration 10 mM and incubated for 60 min at 37 °C. Reactions were terminated with 100 μL of stop solution (50 mM Tris pH 8.0, 20 mM EDTA, and 0.1 ug/mL glycogen) and RNA was precipitated with 500 μL ethanol by incubating for 30 min at −80 °C followed by centrifugation for 30 min at 16,000 g at 4 °C. Supernatant was removed, and pellet resuspended in H2O.

To analyze NAD-capping of DNAzyme-generated fragments of mitochondrial RNAs, 25 μg of cleaved RNA was separated by electrophoresis on 8% urea polyacrylamide gels supplemented with 0.3% 3-acrylamidophenylboronic acid (Boron Molecular). RNA was transferred to positively charged Nylon transfer membrane (GE Healthcare Life Sciences), immobilized by UV crosslinking, and incubated with a ^32^P-labeled oligodeoxyribonucleotide probe complementary to the 5’-end fragments of target RNAs (Supplementary Table S2). The probes were labeled using T4 polynucleotide kinase (New England BioLabs) and [γ-^32^P] ATP (Perkin Elmer). Reaction products were visualized with Amersham Typhoon RGB Biomolecular Imager (GE Healthcare Life Sciences) and bands corresponding to uncapped and NAD- or NADH-capped DNAzymes-generated fragments were quantified with IQ TL image analysis software.

### Isolation of mitochondria and quantification of intramitochondrial NAD levels

Mitochondria were isolated using density gradient centrifugation as previously described ^43^. Seven grams of exponentially grown cells were used to retrieve pure mitochondria from both YPD (fermentation) and YPG (respiration) cultures. Cells were first treated with Zymolyase (Sunrise Science Products) to generate spheroplasts, which were next lysed using a homogenizer. The unbroken cells, nuclei, and cell debris were next pelleted by three consecutive centrifuges – first for 5 min at 1500 g at 4 °C, second at 3000 g for 5 minutes and the third one at 12000 g for 15 minutes. The resulting supernatant was next layered onto a sucrose gradient (15 %-60 %) and purified mitochondria were isolated using density gradient centrifugation (134,000 g for 1 hour). The intact mitochondria forms a brown band at the 60 % −32 % sucrose interface. Mitochondrial preparation purity was assessed by Western blot analysis Anti-Cox2 (Anti-MTCO2 antibody [4B12A5] (ab110271)), and anti-Pgk1 ((PA5-2861) Invitrogen) antibodies were used to detect the respective proteins.

Mitochondrial NAD levels were quantified from three independent mitochondrial preparations from both WT and *xrn1-H41A* strains using NAD/NADH Quantitation Kit (Sigma-Aldrich). Mitochondrial extract corresponding to10 μg of protein from both WT and *xrn1-H41A* were used to determine NAD levels as per instructions provided in the kit.

## SUPPLEMENTARY FIGURES

**Supplementary Figure S1.**
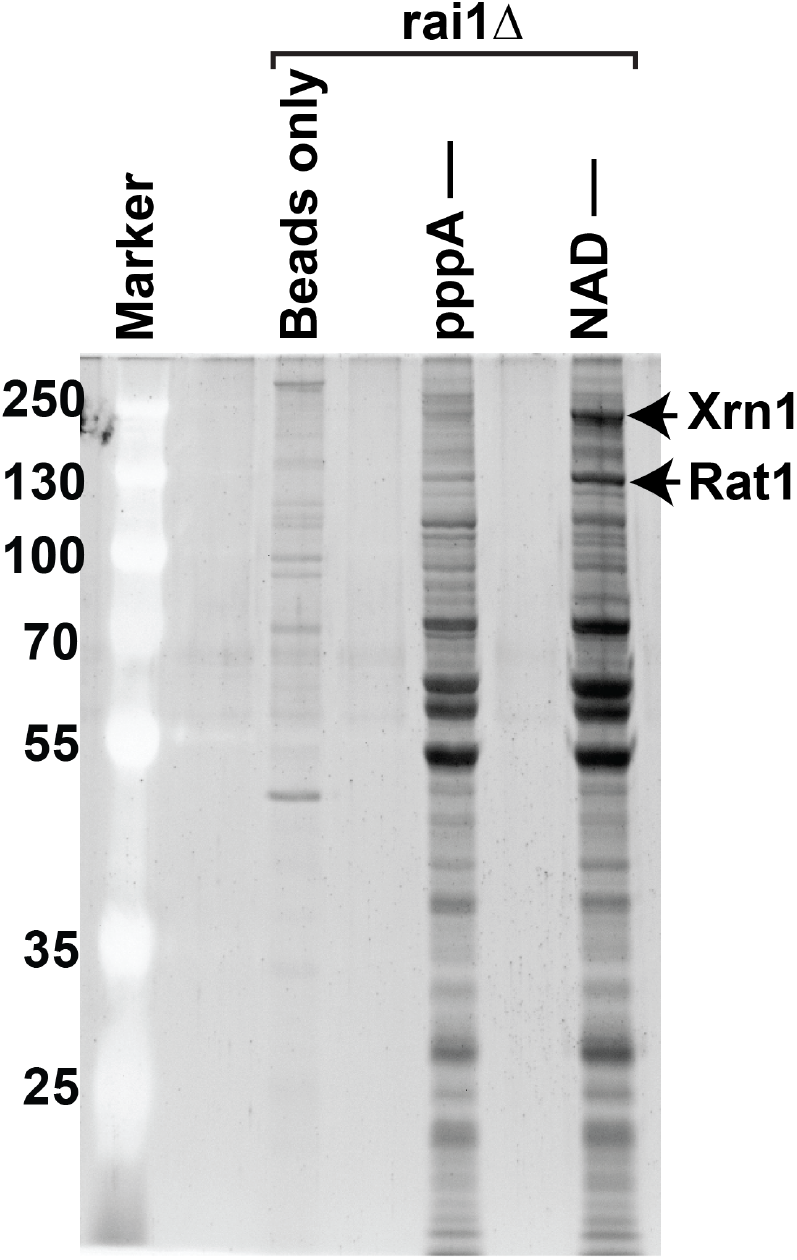
Identification of proteins bound to the NAD cap in the absence of Rai1. Protein derived from a strain lacking Rai1 (*rai1Δ*) were captured by NcRAP and eluates detected by SYPRO Ruby following resolution on 10% SDS-PAGE gel. Affinity purification with 5’-end triphosphorylated RNA (pppA---) or NAD-capped RNA (NAD---) are shown.

**Supplementary Figure S2.**
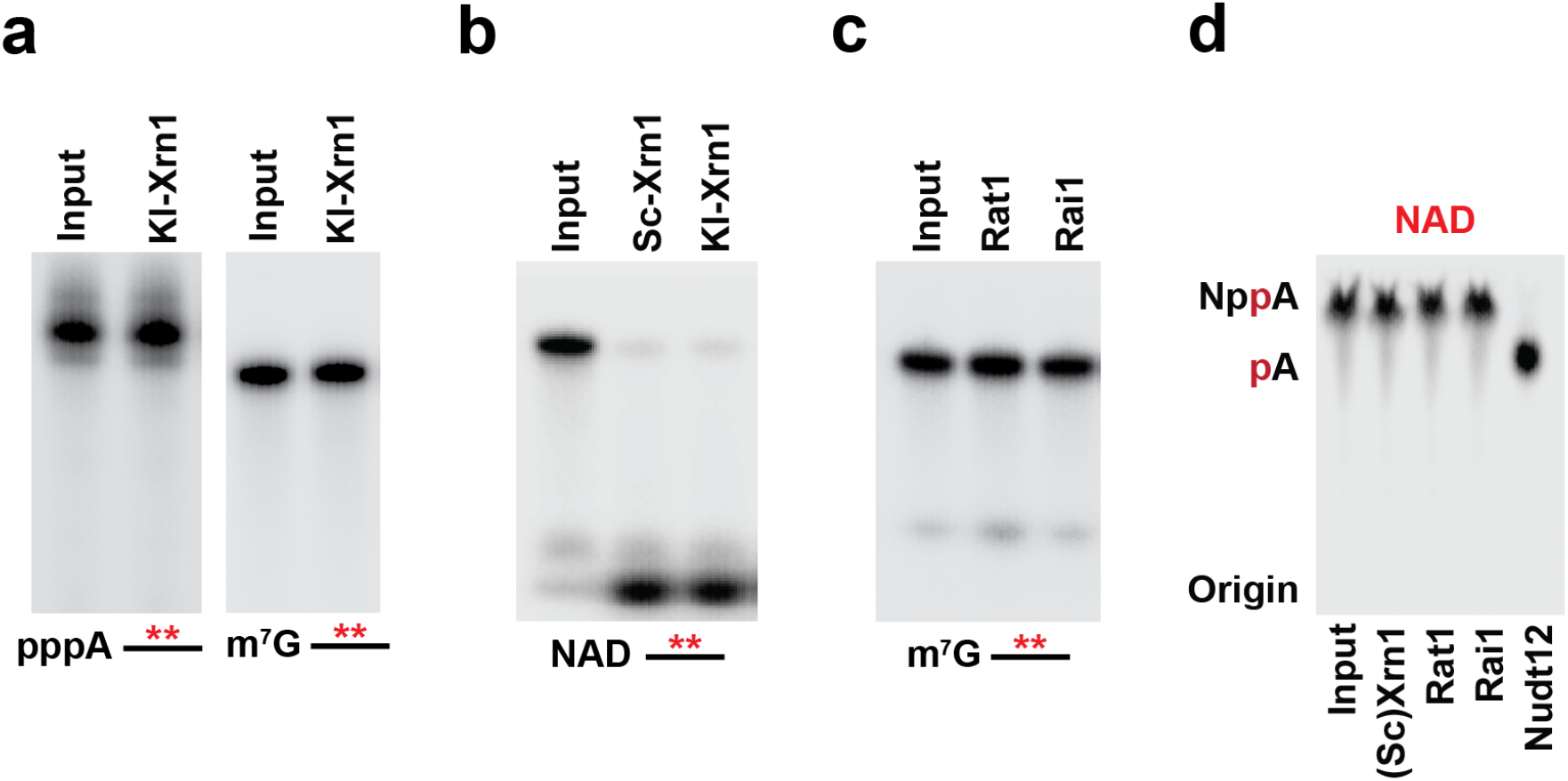
Xrn1 hydrolyzes NAD-capped RNA but not 5-triphosphorylated or m^7^G-capped RNA. (**a**) Recombinant *K. lactis* Xrn1 (30 nM) was incubated with uniformly ^32^P-labeled 5’-triphosphorylated (pppA) or m^7^G-capped RNA. Products were resolved by 15% 7M urea PAGE. **(b)** Recombinant Xrn1 from *K. lactis* or *S.cerevisiae* (30 nM) were incubated with NAD-capped RNA uniformly ^32^P-labeled and the products were resolved as in panel (A). **(c)** *S.pombe* Rat1 (60 nM) or Rai1 (25 nM) were incubated with uniformly ^32^P-labeled RNA containing a 5’-end m^7^G cap and products resolved as in (A). **(d)** ^32^P-labeled free NAD was incubated with *S. cerevisiae* Xrn1 (30nM), *S.pombe* Rat1 (60 nM) or Rai1 (25 nM), or mouse Nudt12 (50 nM). The reaction products were resolved by polyethyleneimine (PEI)-cellulose thin layer chromatography (TLC) developed in 0.45M (NH4)2SO4.

**Supplementary Figure S3.**
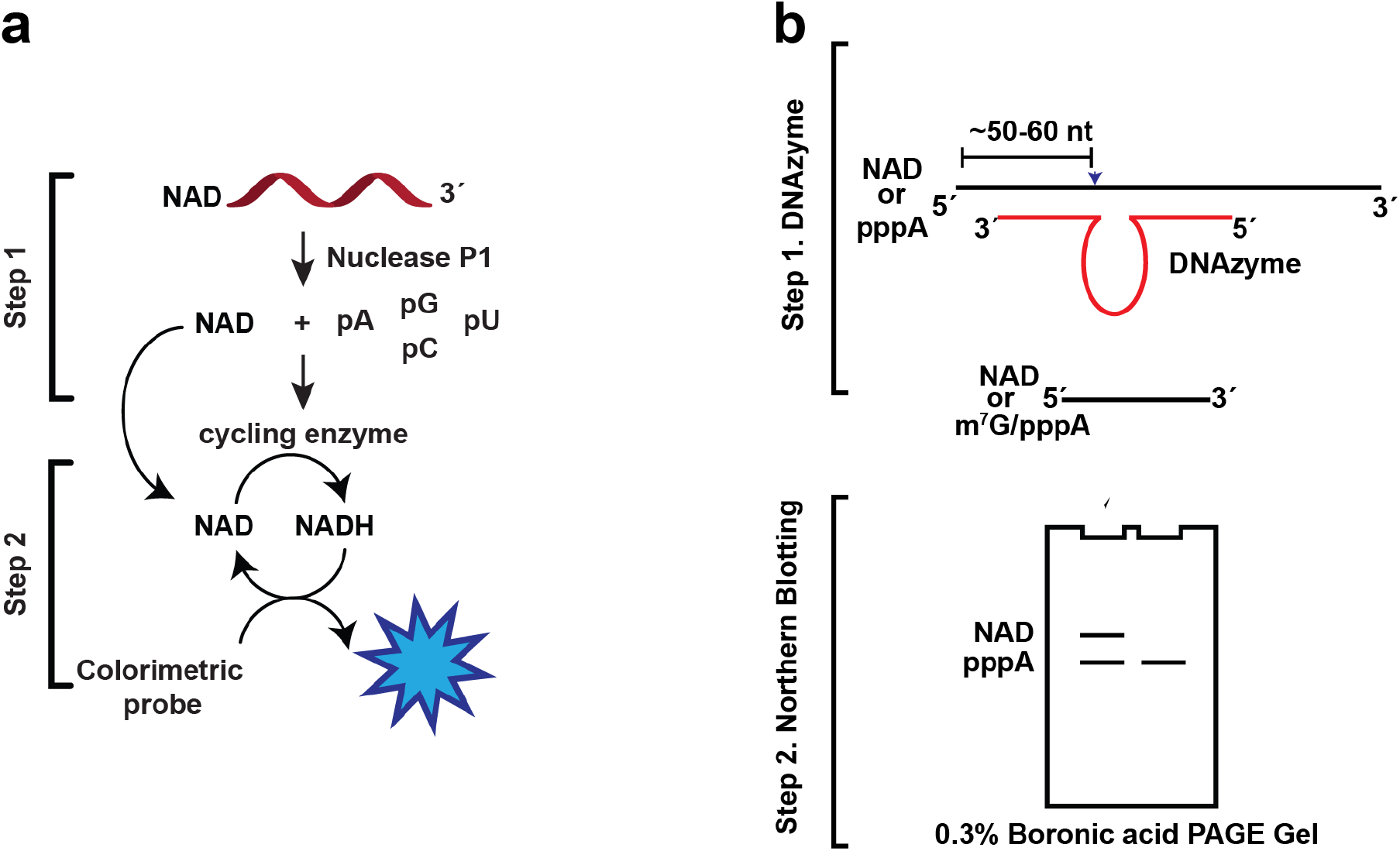
NAD-capQ and DNAzyme. **(a)** Schematic illustration of NAD-capQ as described in ^21^. **(b)** Schematic representation of DNAzyme-mediated RNA cleavage coupled to Northern Blot detection of NAD-capped RNAs.

**Supplementary Figure S4.**
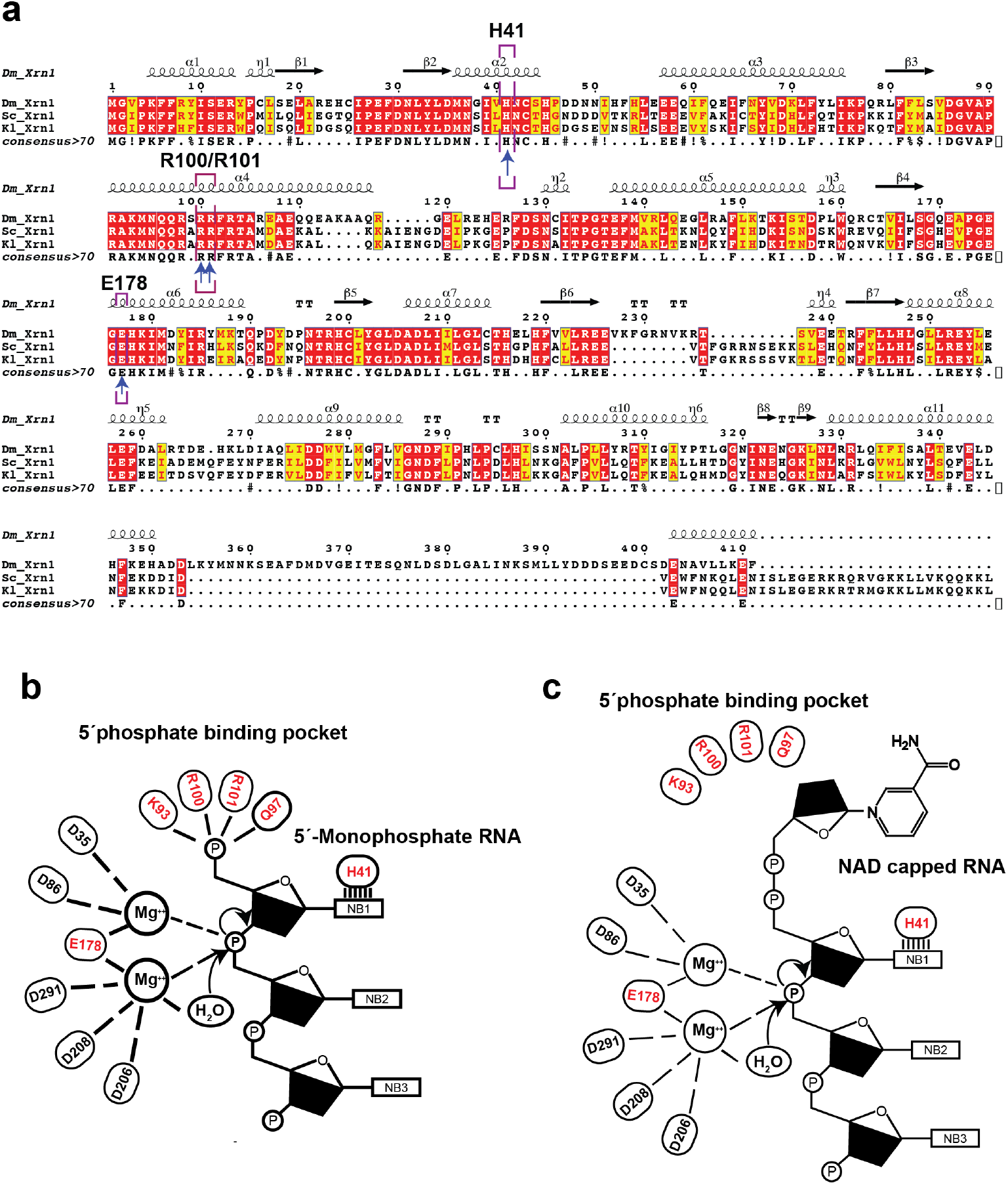
Xrn1 sequence alignment. **(a)** Amino acid sequences corresponding to the amino termini of Xrn1 from *D. melanogaster*, *S. cerevisiae* and *K. lactis* were aligned using ClustalW2 (EMBL-EBI) and the alignment file was analyzed with EsPript 3.2. Amino acid identity is marked in red and similarities are represented in yellow. The catalytically active residues are highlighted with purple rectangles. **(b)** Model for monophosphate RNA hydrolysis (Jinek et al 2012). **(c)** A putative model for processive NAD-capped RNA degradation by Xrn1. According to our *in vitro* and *in vivo* results, the predicted π-π stacking between the His41 and adenosine residue of NAD is critical for the deNADding reaction. This reaction is likely the bottleneck for NAD cap hydrolysis.

**Supplementary Figure S5.**
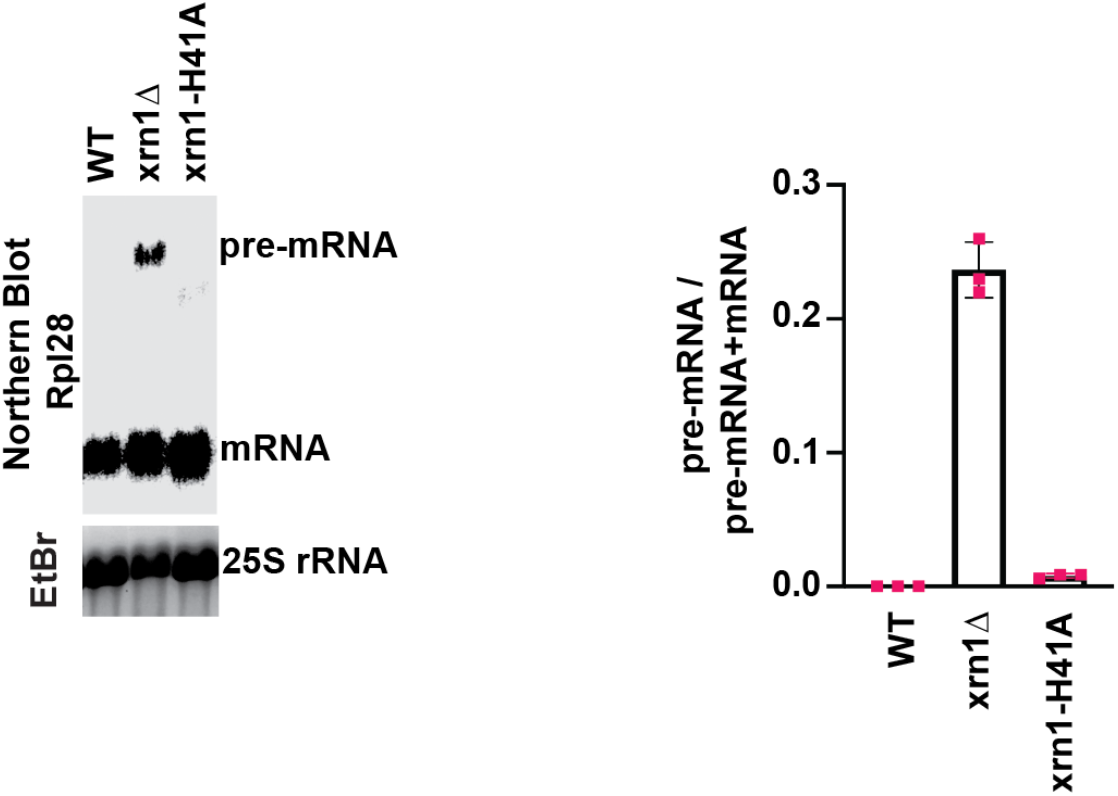
Northern Blot for Rpl28 mRNA in WT, *xrn1Δ* and *xrn1*-H41A knock-in mutant. Quantitation from three independent experiments are presented on the right with the mean ratio of RPL28 pre-mRNA over total (pre-mRNA + mRNA) for the WT, *xrn1Δ* and *xrn1*-H41A. Error bars represent ± SEM.

## Notes

### Competing Interest Statement

The authors have declared no competing interest.

